# Evaluation of Safety and Immunogenicity of an Adjuvanted, TH-1 Skewed, Whole Virion InactivatedSARS-CoV-2 Vaccine - BBV152

**DOI:** 10.1101/2020.09.09.285445

**Authors:** Brunda Ganneru, Harsh Jogdand, Vijaya Kumar Dharam, Narasimha Reddy Molugu, Sai D Prasad, Srinivas Vellimudu, Krishna M Ella, Rajaram Ravikrishnan, Amit Awasthi, Jomy Jose, Panduranga Rao, Deepak Kumar, Raches Ella, Priya Abraham, Pragya Yadav, Gajanan N Sapkal, Anita Shete, Gururaj Rao Desphande, Sreelekshmy Mohandas, Atanu Basu, Nivedita Gupta, Krishna Mohan Vadrevu

## Abstract

We report the development and evaluation of safety and immunogenicity of a whole virion inactivated SARS-COV-2 vaccine (BBV152), adjuvanted with aluminium hydroxide gel (Algel), or a novel TLR7/8 agonist adsorbed Algel. We used a well-characterized SARS-CoV-2 strain and an established vero cell platform to produce large-scale GMP grade highly purified inactivated antigen, BBV152. Product development and manufacturing were carried out in a BSL-3 facility. Immunogenicity was determined at two antigen concentrations (3μg and 6μg), with two different adjuvants, in mice, rats, and rabbits. Our results show that BBV152 vaccine formulations generated significantly high antigen-binding and neutralizing antibody titers, at both concentrations, in all three species with excellent safety profiles. The inactivated vaccine formulation containing TLR7/8 agonist adjuvant-induced Th1 biased antibody responses with elevated IgG2a/IgG1 ratio and increased levels of SARS-CoV-2 specific IFN-γ+ CD4 T lymphocyte response. Our results support further development for Phase I/II clinical trials in humans.

## 1. Introduction

Severe Acute Respiratory Syndrome Coronavirus 2 (SARS-CoV-2), a novel human coronavirus ^1^, has spread to almost every country in the world. SARS-CoV-2 belongs to β-genus of serbecovirus and is a close relative of SARS-CoV with close to 80% sequence identify. The World Health Organization (WHO) declared the disease caused by SARS-CoV-2, Coronavirus Disease-19 (COVID19), a pandemic in March 2020. So far, SARS-CoV-2 has infected more than 25 million people causing close to 850,000 deaths. It is, therefore, imperative to develop effective prophylactic and therapeutic countermeasures to prevent and treat COVID19.

The development of a safe and effective vaccine has become a top priority globally to prevent the spread of SARS-CoV-2 infection during the pandemic. Numerous vaccine candidates are in the preclinical and clinical trial stages. However, meeting the global need for billions of doses of COVID-19 vaccines will require collective effort to identify, evaluate, validate, and manufacture effective vaccines. Inactivated vaccines for viral diseases have been licensed for decades with well-established safety profiles^2^. The availability of well-characterized vero cell manufacturing platform with proven safety in other licensed, live, and inactivated vaccines have aided in rapid vaccine development ^3, 4, 5, 6, 7^. Prior experience in developing inactivated had given us the confidence to develop a fully inactivated with an intact virion, imperative for obtaining an antigen that will yield high immunogenicity. Therefore, to facilitate the development of an effective COVID19 vaccine, we have used a well-characterized SARS-CoV-2 strain and an established vero cell (CCL-81) platform to produce large-scale GMP grade highly purified BBV152 vaccine candidate. It has to be mentioned here that there are several vaccine candidates at different stages of clinical development, such as adenovirus-vectored vaccines, recombinant protein-based, and inactivated vaccines. The inactivated vaccine (PiCoVacc) and the recombinant vaccine (CoV-RBD219N1), which are aluminium adjuvant formulations, have been shown to generate high levels of neutralizing antibodies (NAb) to the S-protein, which could play an important role in vaccine efficacy. Hence, the development of inactivated vaccines for COVID-19 disease prevention appears to be a rational approach, while recognizing the fact that such inactivated vaccines with alum adjuvant specifically induce T helper 2 cells.

While the development of safe and effective coronavirus vaccines is a priority, vaccine-induced disease enhancement observed in preclinical animal models due to Th2-like immunity is a concern. To circumvent the Th-2 bias and to develop a safe vaccine, we formulated a new adjuvant that contains an imidaquizoquinoline class TLR7/8 agonist adsorbed to Algel. TLR7/8 agonists induce strong type I interferon responses from dendritic cells and monocyte-macrophages that facilitate the development of Th1 biased immunity instead of a pathogenic Th2-biased immunity ^8^.

Here, we report the immunogenicity and safety evaluation of the whole virion inactivated SARS-CoV-2 vaccine candidate BBV152, which was evaluated at three antigen concentrations (3,6, and 9μg) and two adjuvants in three animal models, i.e., mice, rats, and rabbits. Our results show that these vaccine formulations induced significantly elevated titers of antigen binding and neutralizing antibodies in all animal models tested without any safety concerns. We also show that the vaccine was formulated with Algel-adsorbed TLR7/8 agonist-induced Th1 biased immunity with significantly elevated SARS-CoV-2 specific IFN⍰+ CD4 T cell response. Collectively these results demonstrate that the BBV152 vaccine candidate induces protective and durable NAb and T cell responses. As a result, BBV152 vaccine candidate has been considered for phase I clinical trials.

## 2. Results

### 2.1 Isolation and selection of SARS-CoV-2 strain for vaccine candidate preparation

During the initial outbreak of SARS-CoV-2 in India, specimens from 12 infected patients were collected and sequenced at the Indian Council of Medical Research-National Institute of Virology (ICMR-NIV), India, a WHO Collaborating Center for Emerging Viral Infections ^9^. The SARS-CoV-2 strain (NIV-2020-770) used in developing the BBV152 vaccine candidate was retrieved from tourists who arrived in New Delhi, India^10, 11^. The sample propagation and virus isolation were performed in the Vero CCL-81. The SARS-CoV-2 sequence was deposited in the GISAID (EPI_ISL_420545). The BBV152 vaccine candidate strain is located in the (G clade), also represented as ‘20A’ clade that is the most prevalent strain in India (followed by ‘19A’) as per data represented in the Next strain analysis of the Indian analysis^12^. In terms of the overall divergence of SARS-CoV-2, this strain is 99.97% identical to the earliest strain Wuhan Hu-1^13^. The multiple passages done in the Vero CCl-81 demonstrated the genetic stability of the virus. The next-generation sequencing (NGS) reads generated from the nucleotide sequences of the BBV152 vaccine candidate strain and its passage one at PID-3 was found to be comparable with the SARS-CoV-2 Wuhan Hu-1 strain (**Table 1**). A maximum difference of 0.075% in the nucleotides was observed, indicating negligible changes in the different batches of the samples analyzed—these results showed genetic stability of the NIV-2020-770 strain for further vaccine development. The seed virus (NIV 2020-770 strain) was transferred from ICMR-NIV to Bharat Biotech, India. Samples from different drug Substance batches of BBIL, along with the original virus, clustered into a single group and indicate an origin from previous passages.

**Table 1:** Genetic Stability of the BBV152 viral strain under specific passages **(Vero CCL-81 Passage 1 PID-3)**

| Reference Position | Wuhan Hu-1 nucleotide | Current nucleotide | Count of reads | Frequency of reads | Region |
| --- | --- | --- | --- | --- | --- |
| 241 | C | T | 10937 | 99.7 | 5' UTR |
| 3037 | C | T | 6227 | 99.6 | orf1ab |
| 4809 | C | T | 11561 | 99.91 | orf1ab |
| 14408 | C | T | 7562 | 99.91 | orf1ab |
| 23403 | A | G | 13336 | 99.96 | S |

### 2.2 Vaccine candidate preparation

GMP production of virus bulk was standardized in bioreactors. The seed virus was adapted to a highly characterized GMP vero cell platform, amplified to produce the master and working virus bank. The master virus bank was characterized based on WHO Technical Report Series guidelines (identity, sterility, mycoplasma, virus titration, adventitious agents, hemadsorption, virus identity by Next Generation Sequencing. The viral RNA isolated from the master virus bank (MVB) was sequenced using NGS the platform at ICMR-NIV and Eurofins Bangalore, India. The sequence reconfirmed the identity of MVB as the NIV 2020-770 strain of SARS-CoV-2.

Vero cells and virus were propagated in the bio-safety level-3 (BSL-3) facility using bioreactors. Growth kinetics analysis showed that the stock replicated to 7.0 log10 TCID_50_ between 36- and 72-hours protection. ß-propiolactone was utilized for the inactivation of the virus by mixing the virus stock between 2-8°C. During the inactivation kinetics experiments with varying conditions and concentrations, samples were collected at various time points (between 0 to 24 hours, at 4-hour intervals) to evaluate the cytopathic effect of live virus. Three consecutive inactivation procedures were performed to ensure complete viral inactivation without affecting the antigen stability **(Figure 1A)**. Transmission electron microscopy (TEM) analysis showed that the inactivated and purified virus particles were intact, oval-shaped, and were accompanied by a crown-like structure representing the well-defined spike on the virus membrane **(Figure 1B)** Inactivated and purified virus was also characterized by western blot for its identity with SARS-CoV-2 specific antibodies using various stages of vaccine candidate development such as cell harvest, clarified supernatant, post-inactivation, and purification. Western blot analysis showed distinct bands of all proteins. Purified and inactivated whole virion antigen produced from three production batches were probed with anti-Spike (S1 & S2), anti-RBD, and anti-N protein **(Figure 1C)**. These results showed that the final purified inactivated bulk of the vaccine candidate is highly pure and contains S (S1, S2), RBD, and N protein bands with their corresponding equivalent molecular weight.

**Figure 1:**
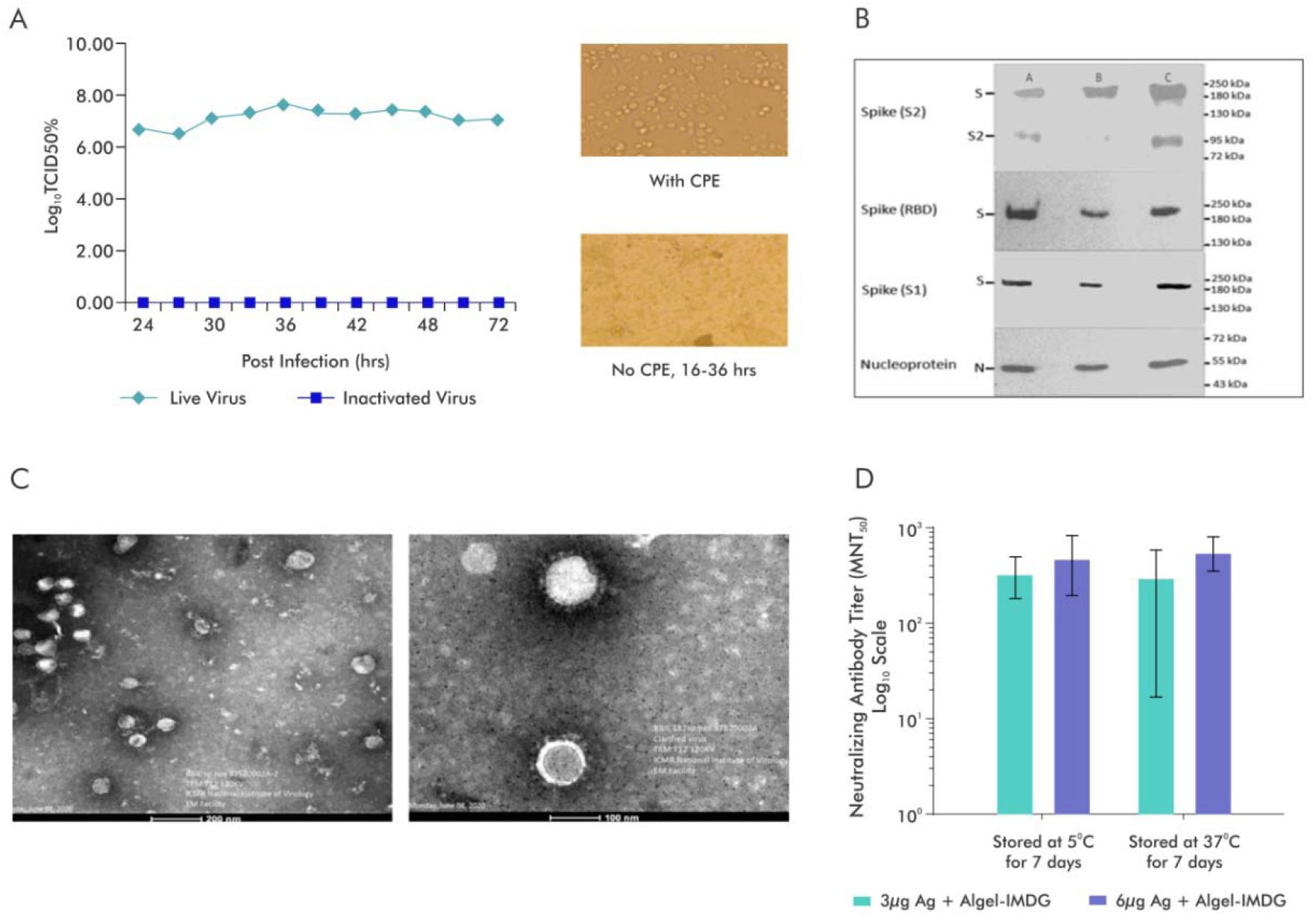
Characterization of inactivated SARS-CoV-2 and evaluation of the stability of BBV152 vaccine formulations. **A**.SARS-CoV-2 Virus (Strain NIV-770-2020) Growth Kinetics & Cytopathic effect (CPE) of virus before and after Inactivation (i) Virus titer (10^6^-10^7^) measured by CCID_50_ at every 3 hours up to 48 and after that every 12hrs various time points (24, 27, 30, 33, 36, 39, 42), (ii) Cells with Cytopathic Effect (CPE) before inactivation and No CPE after Inactivation, (iii) Image of Vero cell monolayer with no CPE observed from 16-36hrs; **B**. Representative electron micrograph of purified inactivated SARS-CoV-2 candidate vaccine (BBV152) at a scale bar: 100 nm (right) and 200 nm (left); **C**. Western blot analysis of Purified Inactivated SARS-CoV-2 produced from three production batches; **D**. Microneutralization antibody titer of Day 14 sera collected from mice vaccinated with Adjuvanted formulations (3μg Ag with Algel-IMDG and 6μg Ag with Algel-IMDG), after subjecting them for stability at 37°C for 7 days and compared with 2-8°C

### 2.3 Vaccine formulations with adjuvants

BBV152 vaccine candidates were formulated with two alum adjuvants: Algel (aluminium hydroxide gel) and Algel-IMDG, an imidazoquinoline class molecule (TLR7/TLR8 agonist abbreviated as IMDG) adsorbed on aluminium hydroxide gel. The agonist molecule for Algel-IMDG was licensed from ViroVax LLC, USA. Three vaccine formulations were prepared with 3μg and 6μg with Algel-IMDG (BBV152A and BBV152B, respectively) and 6μg with Algel (BBV152C). To determine the stability of the vaccine formulations, inactivated antigen plus adjuvant preparations were stored at 37°C and 2-8°Ctemperature for seven days. These vaccine formulations were evaluated in Balb/C mice to estimate Nab titer by microneutralization test (MNT_50_). Our results demonstrated that the vaccine formulations are relatively stable at 37°C for 7 days, as shown by equivalent Nab titer compared to formulation stored at temperature 2-8°C **(Figure 1D)**. There is no significant difference between the two formulations (BBV152A & BBV152B)

### 2.4 Safety

All the three BBV152 formulations, the pure antigens at 3 different concentrations, and the two adjuvants have been evaluated for safety in three animal models (mice, rats, and rabbits) following the required regulatory guidelines^14, 15, 16, 17^. **Table 2** summarizes the key tests completed and the observations thereof. Safety has been established in repeat-dose toxicity studies in Balb/C mice (female, 6-8weeks old) which were vaccinated intraperitoneally (*i.p*) with 1/20^th^of the intended human single dose (HSD, 3 or 6 or 9 μg) of inactivated vaccine candidate with or without adjuvant on day 0, 7 and 14. In contrast, New Zealand white rabbits, Swiss Albino mice, and Wistar rats were vaccinated intramuscularly (*i.m*. Algel-IMDG alone was further evaluated for safety by mutagenicity assay (bacterial reverse mutation). No substantial increase in revertant colony numbers in any of the tested strains was observed following treatment with Algel-IMDG alone at any dose level, in both the plate-incorporation and pre-incubation methods in the presence or absence of metabolic activation (S9 mix). The positive controls (Sodium azide, 4-Nitro-o-phenylenediamine, Methyl methane sulfonate, and 2-Aminoanthracene) used for various strains showed a distinct increase in induced revertant colonies in both the methods. **(Figure S1)**

**Table 2:** Safety studies conducted

| Study Type | Test System | Test Item<sup>1-3</sup> | Route of Administration | Key Test Item result |
| --- | --- | --- | --- | --- |
| Repeated dose toxicity studies | Wistar Rats | Antigen, Adjuvanted vaccines, & Adjuvants | Intramuscular | All the Test Items have been demonstrated to be safe from a Toxicology perspective<sup>4</sup>. |
|  | Swiss albino Mice | Adjuvanted vaccines & Adjuvants | Intramuscular |  |
|  | BALB/c Mice | Antigen, Adjuvanted vaccines, & Adjuvants | Intraperitoneal |  |
|  | New Zealand White Rabbits | Adjuvanted Vaccines | Intramuscular |  |
| Mutagenicity assay (Bacterial Reverse Mutation) | *Salmonella typhimurium* | Algel-IMDG | -- |  |
| Maximum Tolerated Dose studies | Swiss albino mice & Wistar Rats | Algel-IMDG | Intramuscular |  |

1. Antigen: BBV152 Antigen at 3, 6 & 9µg. 2. Adjuvanted vaccines: BBV152A, BBV152B & BBV152C. 3. Adjuvants: Algel & Algel-IMDG at 200 & 300µg. 4. Details are given in Supplementary Section.

In the Maximum Tolerated dose study performed with Algel-IMDG, the test item was tolerated at the tested dose (200 μg/animal) in mice and rats as demonstrated by lack of erythema, edema, or any other macroscopic lesions at the site of injection. Algel is a well-known adjuvant having been used in a large number of vaccines globally, we evaluated the safety profile of the novel adjuvant used in this study. Histopathology examination of the injection site showed active inflammation, as demonstrated by mononuclear cell infiltration, which is likely a physiological local inflammatory reaction caused by aluminium salt in the vaccine adjuvant preparation. In any of the studies conducted, there were no mortality or no changes observed in clinical signs, body weight gain **(Figure S2)**, body temperature, or feed consumption in the treated animals.

### 2.5 Clinical pathology investigations

In all the animal models, haematology, clinical biochemistry, coagulation parameters, and urinalysis treated with adjuvanted vaccine candidates or adjuvants/ antigen alone were comparable to control **(Figure S3)**. The following exceptions were noticed as Alpha 1-acid glycoprotein values were increased on day 2 with Algel-IMDG in male rats when compared to day 0, which reduced to normal levels by day 21. Evidence of an acute phase response was indicative of reactogenicity to the vaccine formulation, and the increase was noticed in the adjuvanted vaccine with the Algel-IMDG group alone. These findings correlate with the inflammatory reaction at the injection site in this group. The absolute and relative neutrophil counts were increased in female rats of groups (Antigen 6 μg + Algel 300 μg), and (Antigen 9 μg + Algel-IMDG 300 μg) on day 2 as compared to control. However, these values were noticed and were comparable to control on Day 21. This transient increase may be due to inflammation at the injection site after administration of the first dose.

### 2.5 Necropsy, organ weight, and histopathology

There were no treatment-related microscopic findings observed in antigen alone by intramuscular (*i.m*) route. In groups treated with adjuvants alone and adjuvanted vaccine with Algel &Algel-IMDG, local reaction at the site of injection (quadriceps muscles of the hindlimb) was observed. In animals treated with Algel alone or adjuvanted vaccine with Algel, inflammatory changes characterized by mild infiltration of mononuclear cells and the presence of macrophages containing bluish material (interpreted to be aluminium in Algel) were observed. On day 21, animals treated with Algel-IMDG alone showed inflammation around homogeneous bluish material (interpreted to be test item) characterized by the infiltration of mononuclear cells. Additionally, macrophages containing bluish stained material understood to be aluminium in the test item (Algel-IMDG) were also observed. Algel produced a milder reaction when compared to Algel-IMDG. On day 28, reduction of inflammation was observed in both the adjuvants, and the number of macrophages containing bluish stained material was also observed less in the recovery groups when compared to day 21 **(Figure S4 & S5)**. No microscopic findings were observed in any of the organs examined, including spleen and lymph nodes, any of the animal models **(Figure S6 & S7)**. Organ weights across groups were comparable.

### 2.6 Immunogenicity studies

We assessed the immunogenicity of BBV152 formulations in BALB/c mice and New Zealand white rabbits]. All immunization studies were conducted based on a three-dose IM regimen conducted on days 0, 7, and 14. Pooled or individual serum samples collected on days 0, 7, 14, and 21 post-immunization/boost were evaluated for antibody binding (ELISA) and Nab by plaque reduction neutralizing titer (PRNT_90)_or MNT_50_against live SARS-CoV-2 strain.

### Immunogenicity inBAlb/c Mice

To assess the immunogenicity of the candidate vaccines, BALB/c mice (n=10) were injected via i.*p* route with three concentrations of antigen at 1/20^th^of the intended human single dose (i.e., 3μg, 6μg, and 9 μg/mouse). Vaccine formulations adjuvanted tested at three antigen concentrations elicited high levels of binding and Nab titer **(Figure2 A & B)**. Antigen alone and adjuvants alone were included in these studies as controls (data not presented for brevity). Further, to assess the immunogenicity and safety of clinical batch samples, Balb/C mice (n=10/group, 5 Male and 5 Female) were vaccinated via IP route with three adjuvanted formulations with Algel and Algel-IMDG at 1/10^th^ human intended single dose (3, and 6 μg/dose with Algel or Algel-IMDG). All adjuvanted vaccine formulations elicited antigen-specific binding antibodies **(Figure 2C)**. Further, sera collected on Day 21 were analyzed by ELISA to determine S1, RBD, and N specific binding titer **(Figure 2D)**. Analysis of PRNT_90,_ performed with individual mice sera, showed high Nab’s in all adjuvanted vaccines **(Figure 2E)**, while **Figure 2F** depicts the effect of dose sparing of Algel-IMDG. Figure S8 depicts the 8-fold increase in vaccine potency, when dosing was in day 14 intervals. Additionally, dosing with antigen alone was found to be immunogenic. However, the responses were significantly lower than the adjuvanted vaccine **(Figure S9)**.

**Figure 2:**
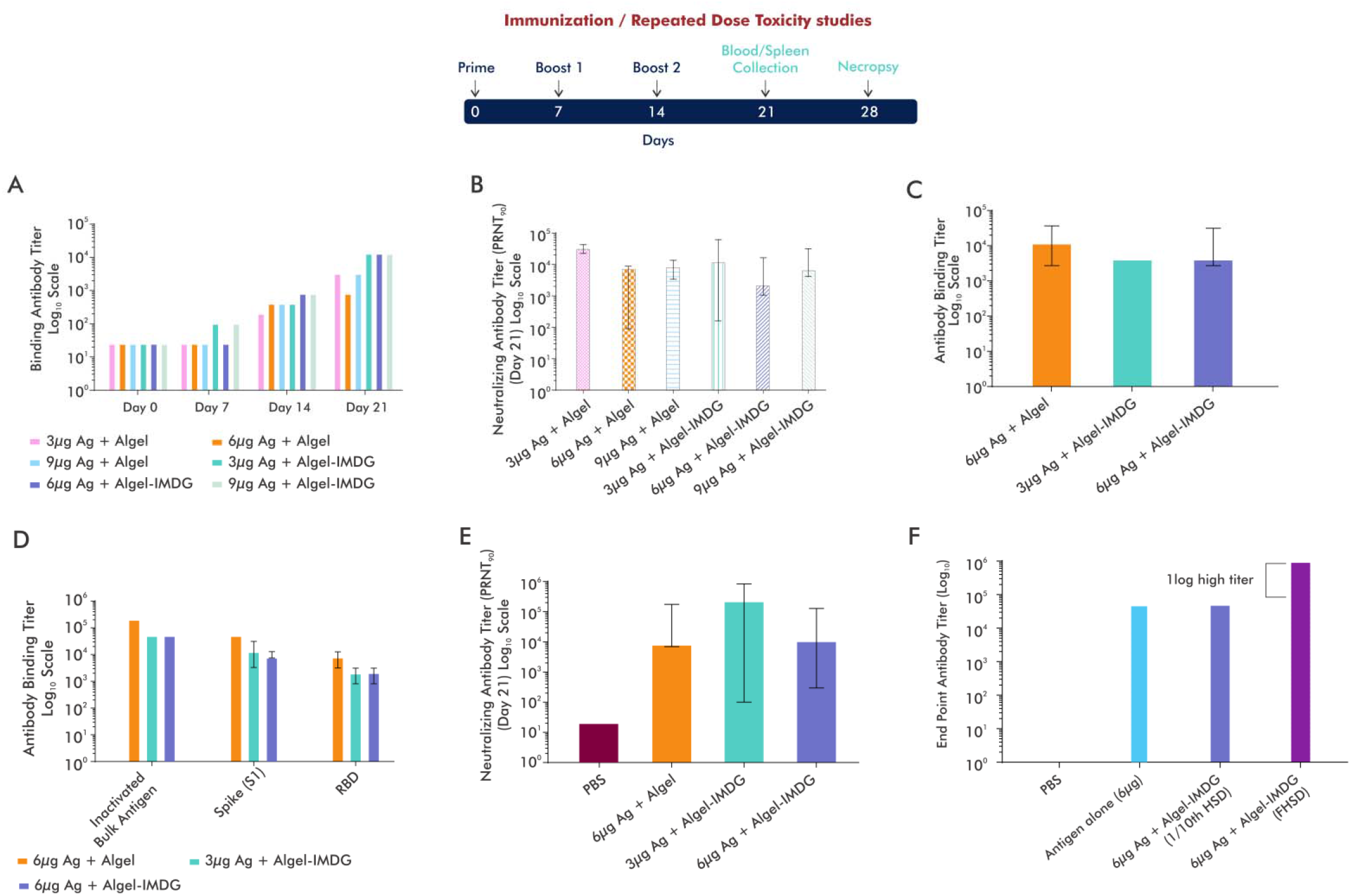
BBV152 Vaccines Induces High Virus-specific Antibody Response in Mice and Rats. Balb/C mice (n=10) were administered with adjuvanted vaccine formulations via IP route either with 1/20^th^ (Fig A&B) or 1/10^th^ (Fig C, D & E) Human Single Dose (HSD): **A**. S1 specific Total IgG antibody binding titer performed by ELISA, using sera collected at various time points (Day 0, 7, 14 & 21); **B**. Neutralizing antibody titers performed by PRNT_90,_ using day 21 sera, when administered with 1/20^th^ HSD respectively; **C**. S1 specific Total IgG antibody binding titer performed by ELISA, using Day 21 sera when administered with 1/10th HSD; **D**. SARS-CoV-2 specific (S1, RBD, N and total inactivated antigen) antibody binding titers elicited against adjuvant vaccines (BBV152A, B & C); **E**. neutralizing antibody titers performed by PRNT90, using day 21 sera, when administered with 1/10th HSD respectively; **F**. Balb/C mice were administered with Antigen & BBV152B via IM route at the specified doses on day 0 & 14. Sera were collected on Day 28 (post 2nd dose) and determined S1 specific antibody titer by ELISA.

### Immunogenicity in New Zealand White Rabbits

Rabbits (n=8) were immunized with antigen concentrations for humans (3 and 6 μg/dose) on days 0, 7, and 14. The groups that received BBV152A & B showed a slightly higher binding antibody response compared to BBV152C **(Figure 3A)**, though not statistically significant. Examination of neutralizing antibody titers revealed high PRNT_90_titers on day 21 are also reported **(Figure 3B)**. Further, NAb’s performed by MNT_50_ were compared with Nabs from a panel of human convalescent sera from recovered symptomaticCOVID-19 patients. **(Figure 3C)**.

**Figure 3:**
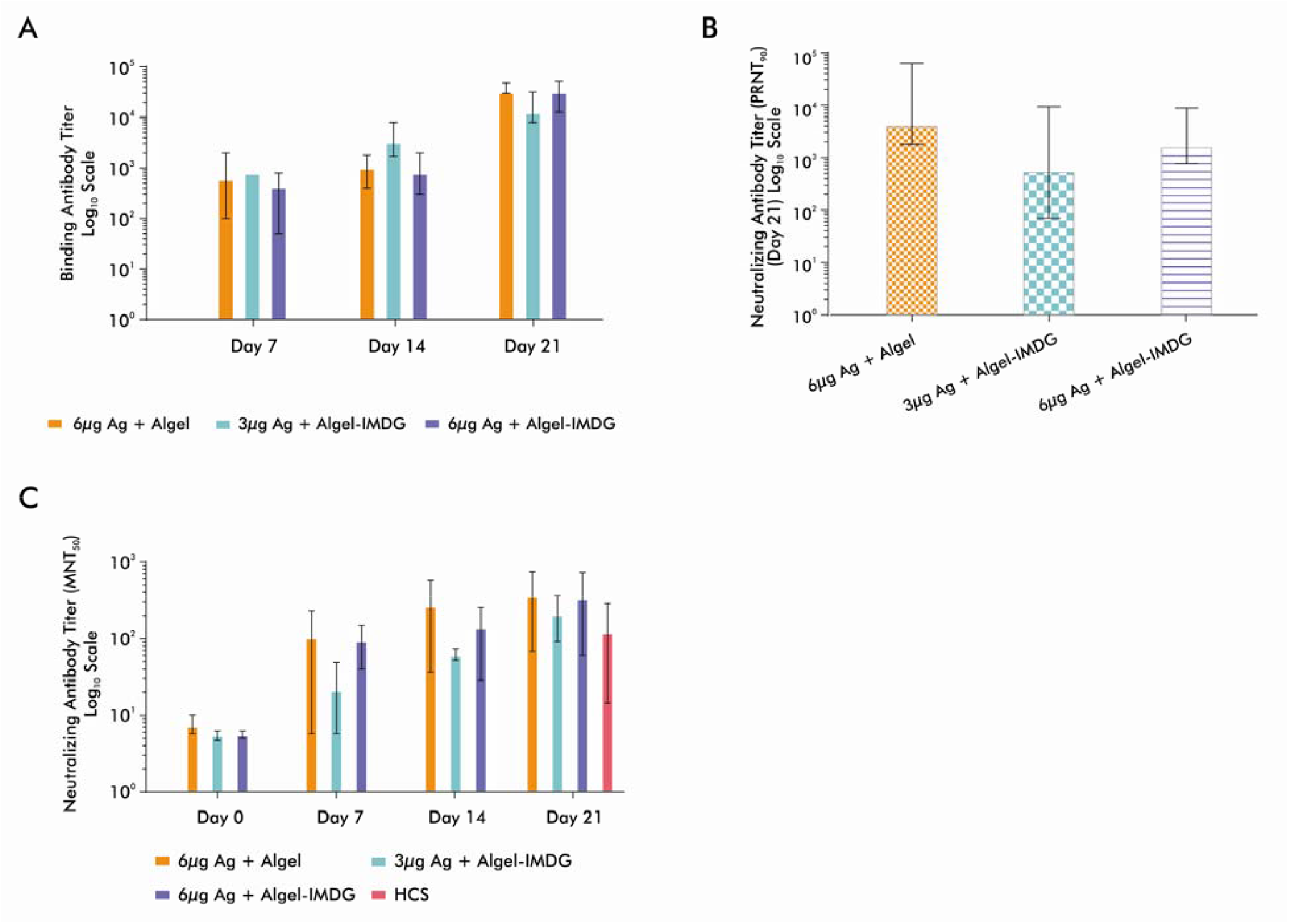
BBV152 Induces Robust Neutralizing Antibody Response in Rabbits. New Zealand white rabbits (n=8) were Rabbits were vaccinated intramuscularly on days 0, 7, and 14 with Full HSD via IM route. SARS-CoV-2 specific Antibody titers were measured by ELISA. Nab tires were measured by PRNT_90_ and MNT_50._ Data Points represent mean ± SEM of individual animal data. A. S1 specific Ab binding titer of sera collected at various time points (Day 0, 7, 14 & 21); B. PRNT_90_ neutralizing antibody titers of Day 21 sera; C. MNT_50_ neutralizing antibody titers of sera collected at various time points (Day 0, 7, 14 & 21) along Neutralizing antibody titer (MNT_50_) with Human convalescent sera (HCS) from recovered COVID-19 patients (n=15). Samples were collected between 21-65 days of virological confirmation.

### BBV152 adjuvanted with TLR7/8 adsorbed algel induces Th1 biased immune response

#### Immunoglobulin Subclasses

Antibody isotyping (IgG1 & IgG2a) was analyzed on day 21 serum samples to evaluate the Th1/Th2 polarization by vaccination with the two adjuvants. The average ratio of IgG2a/IgG1 was higher in all antigen concentrations with Algel-IMDG when compared to Algel, indicative of Th1 bias **(Figure 4A)**. Additionally, antigen immunized with 6μg Algel-IMDG samples induced significantly higher responses of interferon-γ(IFNγ) **(Figure 4B)**. These results suggest that Algel-IMDG adjuvant that contains TLR7/8 agonist induces Th1 biased protective immunity and thus is a promising adjuvant for further development.

**Figure 4:**
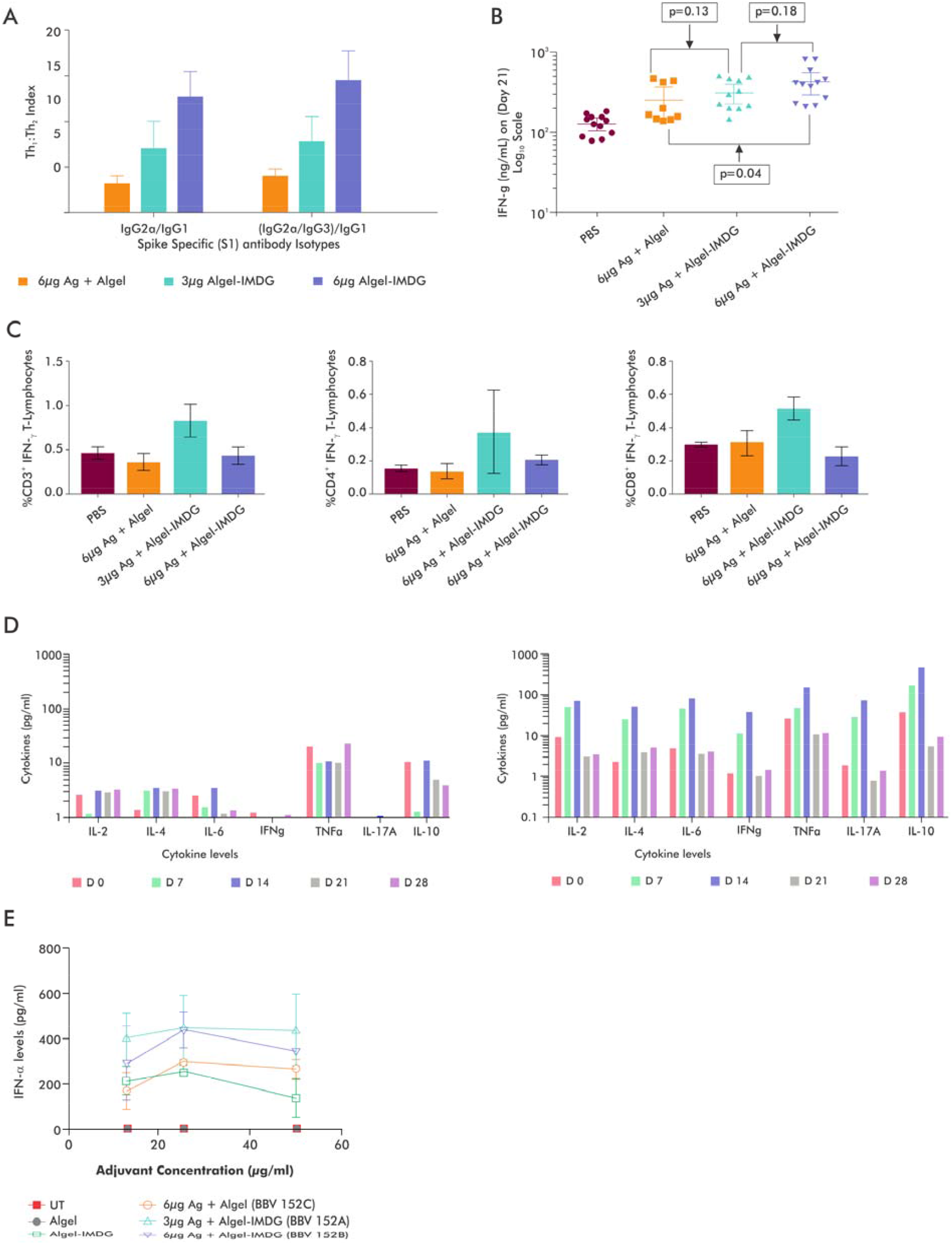
BBV152 Induces A Robust Virus-specific T Cell Response. Balb/C mice (n=10) were vaccinated with 1/10th HSD of adjuvanted vaccine formulations (BBV 152 A, B & C) via the IP route. A. Immunoglobulin subclasses (IgG1, IgG2a & IgG3) were measured by ELISA. Th1:Th2 index was measured using the formulas IgG2a/ IgG1 or (IgG2a+igG3)/IgG1. B. IFNgamma estimation by ELISA, on Day 21 sera (7 days post 3rd Dose). Statistical analysis was done Graph Pad Prism version.7.0; C. Bar graph represents mean data of percent CD3^+^ or CD4^+^ or CD8^+^ T lymphocytes producing IFNgamma from the respective group of animals (i) CD3^+^ T lymphocytes population, (ii) CD4^+^ T lymphocytes population, (iii) CD8^+^ T lymphocyte population. Error bars indicate Mean±SD. Vaccinated mice splenocytes from Balb/C mice (n=10), administered with 1/10^th^ HSD via IM route were used for the analysis; D. Cytokine profile measured on various time points using vaccinated Balb/C mice sera, when administered with Adjuvanted vaccine formulations (1/20^th^ HSD via IP route) Left - BBV152C-Antigen 6μg+Algel); Right – BBV152B-Antigen 6μg+Algel-IMDG, E. IFNα levels measured by ELISA from culture supernatant, when treated healthy PBMCs with Algel or Algel-IMDG or adjuvanted vaccine formulations (BBV152A, B & C). Two-fold serial dilutions of the human intended dose of adjuvanted vaccine formulations were used. Corresponding antigen or adjuvant alone concentration were also maintained simultaneously as controls. Error bars indicate Mean±SD of triplicate values.

To further evaluate whether adjuvanted vaccine formulations (with Algel & Algel-IMDG) induced Th1 response or not, we performed intracellular staining using vaccinated mice splenocytes after stimulation with inactivated SARS-CoV-2 antigen and determined IFNγ producing T lymphocytes. Interestingly, we found that the adjuvanted formulation with Algel-IMDG (BBV152A & B) showed elevated levels of IFNγ producing CD4 cell population, compared to those with Algel. These results indicate that antigen formulated with Algel-IMDG skewed towards Th1 mediated response **(Figure 4C)** and induced strong T cell immunity.

### Cytometric Bead Array (CBA)

Expression of TNF**-**αand interleukins was noticeably expressed in the 6μg Algel-IMDG when compared to 6μg Algel **(Figure 4D)**.

**IFNα responses as a function of innate immunity activation** to assess the effect of adjuvants (Algel or Algel-IMDG) on antigen and understanding the critical role of IFNα in both anti-viral and pro-inflammatory cytokine functions, and linking innate immunity and adaptive immunity, we used PBMCs from healthy volunteers to stimulate using the antigen and adjuvanted vaccines for 36-72hrs at both 3 and 6μg antigen concentration, and measured IFNα. We found that the Inactivated antigen itself stimulated Anti-viral Cytokine (IFN-α), an indicator of the first line of defense. Algel-IMDG containing TLR7/8 agonists also stimulated IFN-α & but not the Algel alone. The addition of Algel and Algel-IMDG showed a synergistic effect on Antigen, which was demonstrated by the elevated of IFN-αlevels in the cell supernatant **(Figure 4E)**; the latter adjuvant being more effective.

## 3. Discussion

Here, we report the development of a whole virion inactivated SARS-CoV-2 vaccine candidate (BBV152). The strain used for this candidate is pathogenic in humans and has shown extensive genetic stability and appropriate growth characteristics for the selection of a vaccine candidate. Preclinical toxicity and safety evaluation of the three formulations showed minimal to no adverse events. Our results show that the vaccine formulations induced significantly elevated antigen-binding antibody and Nab responses in the animals immunized, with a distinct Th1 bias observed with Algel-IMDG adjuvanted vaccines. Although the neutralizing antibody titers are not statistically different between the antigen concentration (3μg and 6μg) or the nature of adjuvant, all the formulations tested have exhibited excellent immunogenicity. Our potency results compare quite favorably with those reported in the literature for similar COVID-19 vaccines. Inactivated SARS-CoV-2 vaccine candidate (BBIBP-CorV) has been shown to induce high levels of Nab titers in mice and rats to provide protection against SARS-CoV-2^3^. A purified inactivated SARS-CoV-2 virus vaccine candidate (PiCoVacc) has also been shown to induce SARS-CoV-2-specific NAb in mice and rats. These antibodies potently neutralized 10 representative SARS-CoV-2 strains, indicative of a possible broader neutralizing ability against SARS-CoV-2 strains circulating worldwide^4^.

The risk of antibody-dependent enhancement (ADE) is a serious concern for COVID-19 vaccine development^18, 19, 20, 21^. A few animal studies from animal SARS-CoV-1 and MERS-CoV inactivated or vectored vaccines adjuvanted with alum have shown correlation to Th2 responses resulting in eosinophilic infiltration in the lungs ^18, 19, 20^. Alum is the most frequently used vaccine adjuvant with an extensive safety record. It is desired to have a COVID-19 vaccine that can generate both humoral and cell-mediated immune responses. The response generated from alum is primarily Th2-biased with the induction of strong humoral responses via neutralizing antibodies ^22^. It is not clear if alum alone can stimulate T-cell responses. Complicating adverse events may be associated with the induction of weakly or non-neutralizing antibodies that lead to antibody-dependent enhancement (ADE) or enhanced respiratory disease (ERD), thus warranting COVID-19 vaccines to induce CD4 Th1(interferon-γ, interleukin-2, tumor necrosis factorα) response with minimal Th2 response^23, 24^. Preclinical studies in mice reported that inactivated vaccine-induced eosinophil immunopathology in the lungs upon SARS-CoV infection ^25^ could be avoided using TLR agonist as or in adjuvant formulations. Although current understanding of the risk of COVID-19 vaccine-associated ADE/ERD is limited, the use of TLR7/8 agonists in an adjuvant in SARS-CoV-2 vaccine formulation will minimize Th2 response, if any.

Over many decades it has shown that vaccination is generally a safe and well-tolerated procedure. Nevertheless, toxic actions of vaccines can result from any of the following, drug substance and drug product, including excipients used for formulation. The current preclinical studies conducted with BBV152, adjuvanted with the two adjuvants, did not indicate any undesirable pathological changes and systemic toxicity. Local reactogenicity to adjuvants used in vaccine formulation were the only findings noted. Algel (Alum) is the most commonly used agent as an adjuvant. It has been shown to act by depot formation at the site of injection, allowing for a slow release of antigen. Further, it converts soluble antigens into particulate forms, which are readily phagocytosed^26^. The microscopic findings at the site of injection in the present studies showed the infiltration of macrophages and mononuclear cells. The other adjuvant, Algel-IMDG, contained TLR7/8 in addition to Algel, which was added to augment innate and adaptive immunity, induced slightly higher reactogenicity. IM injection induces a depot effect followed by the passive trafficking of algel particles via lymphatic flow from the interstitial space to the draining lymph nodes, as revealed by IFN-**β**/luciferase reporter mice (unpublished). The lymph node-targeting of Algel-IMDG ensures high adjuvant activity in the target organ (lymph nodes) by enabling the induction of a strong, specific, adaptive immune response while minimizing systemic exposure. The local reaction in the studies conducted was consistent with those available in the literature for these adjuvants, which is a physiological reaction to injection rather than any adverse event^26, 27^.

Collectively, both the adjuvanted vaccines (with Algel and Algel-IMDG), Antigen and Adjuvant alone did not reveal any treatment-related findings, except local reactions when administered through the human intended route (intramuscular) on days 0, 7, and 14 (n+1) with full Human single dose (HSD) or higher than HSD in rodents and non-rodents, thereby establishing the vaccine safety. In our preclinical studies, we demonstrated that all the three inactivated whole virion SARS-CoV-2 vaccine candidates showed 100% seroconversion with high titers of antigen binding and neutralizing antibody responses. Further, the adjuvanted formulation, BBV152B, when immunized in Balb/C mice, showed 10 times higher dose sparing effect compared to antigen alone **(Figure 2F)**. Moreover, these formulations induced immunity that is biased towards Th1 mediated response, as demonstrated by the ratio between IgG2a and IgG1 (greater than 1) **(Figure 4A)**. Additionally, secretion of anti-viral cytokines such as IL-2, IL-4, IL-6, IL-10, IL-17, TNF-alpha, IFNγ was observed on days 7 and 14(7 days after the 1^st^&2^nd^ dose) of vaccination with Algel-IMDG adjuvanted formulations **(Figure 4D)**. Further, the tendency to secrete anti-viral cytokines, IFN-alpha **(Figure 4E)**, might contribute to the activation of the first line of defense mechanisms, which lead to enhanced activation of antigen-presenting cells, such as dendritic cells or macrophages^28, 29, 30^. It is reported that TLR recognition in innate cell population drives early type I IFN production, thereby promotes viral clearance and the early production of proinflammatory cytokines^31, 32^. Though the mechanism of action is yet to be investigated, we hypothesize that this elevated production of IFNα in the Algel-IMDG based Adjuvanted vaccine may provide better protection in the Hamster and NHP homologous challenge study with SARS-CoV-2 virus.

A combination of high neutralizing antibody titers elicited against inactivated antigen alone and the presence intact spike protein on the surface of the virus confirms that the antigen is in the right confirmation and can itself may act as a Th1 inducer with its surface glycoproteins, intracellular viral proteins.

A major limitation of this paper is the lack of protective efficacy results conferred from BBV 152. Additional live challenge studies in hamsters and non-human primates are completed at NIV, India, and results will be published shortly. With no established correlate of protection, we also evaluated human convalescent sera from recovered symptomatic SARS-CoV-2 patients. Samples were collected 21 days after virological confirmation **(Figure3C)**. Furthermore, two other SARS-CoV-2 inactivated vaccines (BBIBP-CorV and PiCoVacc) from China have entered late-stage human clinical trials with published data on the preclinical immune response. Results from these candidates have reported comparable findings, albeit PRNT_50_^33, 34^.

Bharat Biotech has developed a promising inactivated whole virion vaccine candidate which has now entered phase 1/2 clinical development (NCT04471519). The study is designed to evaluate the safety, reactogenicity, tolerability, and immunogenicity of two intramuscular doses of BBV152 in healthy volunteers.

## Methods

### 1. Cells and Virus

Vero CCL-81 (ATCC# CCL 81) cells were maintained in DMEM supplemented with 10% heat-inactivated fetal bovine serum. Vero cells were revived from GMP master cell bank, which was extensively characterized at BioReliance, USA. SARS-CoV-2 (Strain No#NIV-2020-770) was obtained from the National Institute of Virology, a WHO Collaborating Center for Emerging Viral Infections^9^, Pune, India. SARS-CoV-2 strain (NIV-2020-770) sequence was deposited in the GISAID (EPI_ISL_420545).

Specimens from 12 infected patients were collected during the initial outbreak of SARS-CoV-2 at the National Institute of Virology (NIV), India, a WHO Collaborating Center for Emerging Viral Infections^9^. SARS-CoV-2 strain (NIV-2020-770) was passaged in vero cell lines (Vero CCL-81) and sequenced, and the sequence was deposited in the GISAID (EPI_ISL_420545).

### 2. TCID50

The SARS-CoV-2 virus titer was determined by a cytopathic effect (CPE) method assay. Vero cells ATCC-81 (0.2 x 10^6^ cells/mL) were seeded in 96 well plates and incubated for 16-24 hours at 37 °C. Serial 10-fold dilutions of virus-containing samples were added to 96-well culture plate and cultured for 5-7 days in 5% CO_2_ incubator at 37°C, and cells were observed for cytopathic effect (CPE) under a microscope. The virus titer was calculated by the Spearman Karber method ^35^.

### 3. Virus Inactivation

SARS-CoV-2 Virus (NIV-2020-770) was inactivated with β-propiolactone at a ratio ranging from 1:1500 to 1: 3000 at 2-8°C for 24-32 hours and purified by chromatographic purification method. To ensure the effectiveness of the virus inactivation procedure inactivated SARS-CoV-2 virus was inoculated onto vero CCL-81 monolayers and incubated at 37 °C in a 5% CO_2_ incubator and monitored daily for CPE, consecutively for three passages. Further, to reverify the absence of CPE due to supernatant, neat and 10fold dilution of supernatant was inoculated onto Vero cell monolayer and cultured in a 37°C incubator for 5-7 days, and cells were observed for CPE under a microscope.

### 4. Quantitative Real-Time Reverse Transcription Polymerase Chain Reaction (qRT-PCR)

Total RNA was extracted from the virus sample with a QIAamp Viral RNA mini kit (QIAGEN). SARS-CoV-2 RdRP-2 gene primer probes sequences are as follows: RdRP_SARSr-F2-GTGARATGGTCATGTGTGGCGG,R1-CARATGTTAAASACACTATTAGCATA,P2-FAM CAGGTGGAACCTCATCAGGAGATGC-BHQ1. The SARS-CoV-2 reaction was set up containing a master mix of 10 μL (Thermo) and RNA template 10 μL.qRT-PCR was performed under the following reaction conditions: RT step-42°C for 30 min for reverse transcription, Initial Denaturation step: 95°C for 3 min and then 45 cycles of Denaturation95°C for 15 seconds, annealing58°C for 30 seconds - data acquisition, Extension72°C for 15 seconds. Reactions were set on Biorad-CFX96 as per the manufacturers’ instructions.

### 5. Western blotting

Protein samples (∼30 mg) derived from drug substance estimated by Lowry method ^36^ and standard procedures for western blot were adopted. The primary antibodies used were anti-N protein rabbit monoclonal Ab (1:1000 dilution) and anti-S1 or S2 or RBD protein rabbit polyclonal Ab (1:1000 dilution), either sourced from commercial or in-house and human convalescent sera from patients (1:500 dilution) at 4°C. The secondary antibodies goat anti-rabbit IgG H&L (HRP) (GE NA934,1:4000) and HRP-labeled goat anti-human IgG (gamma chain) cross-adsorbed secondary antibody (Invitrogen, 62-8420) (1:1000). Protein bands were visualized in enhanced chemiluminescence (Azure biomolecular imager, USA).

### 6. Formulations Preparation

In the first formulation, BBV152A, 3μg of antigen was mixed with Algel-IMDG, while BBV152B had 6μg of antigen with the same adjuvant (Algel-IMDG), and the third formulation, BBV152C, had 6μg of antigen adsorbed on alum (Algel). Total protein/unbound protein was estimated by the Lowry method^36^.

### 7. Animal husbandry practices

All animal experiments were performed after obtaining necessary approvals from the Institutional Animal Ethics Committee (IAEC). The experimental protocols adhered to guidelines of the Committee for the Purpose of Control and Supervision of Experiments on Animals (CPCSEA) and also as per the Organization for Economic Co-operation and Development (OECD) Principles of Good Laboratory Practice (1997) ENV/MC/CHEM (98)17.

### 8. Immunization

Three animal models were used to evaluate the immunogenicity and safety of the three inactivated whole virion vaccine formulations (BBV152 A, B & C).

#### Mice

Balb/C or Swiss Albino mice (6-8week old) were vaccinated via an intraperitoneal or intramuscular route with either 1/10^th^ or 1/20^th^of full human single dose (BBV152 A, B or C) of inactivated vaccine with or without adjuvant on day 0, 7 & 14 days(n+1 (one extra dose compared to the intended human regimen doses). A formulation with 9μg was also tested.

#### Rats

Wistar Rats (6-8weeks old) were vaccinated intramuscularly with 9μgof inactivated whole virion vaccine with Algel-1 or Algel-2 on days 0, 7 & 14 days (n+1 doses).

#### Rabbits

Zealand white rabbits (3-4 months old) were vaccinated via an intramuscular route with full Human intended single dose (BBV152 A, B or C; n+1 doses). The animals treated were observed up to 14 days, post third dose.

Further, mice and rats were also administered via an intradermal route with full Human intended single dose (HSD, 1.2μg), and rabbits administered full Human intended single dose (HSD, 2.4μg) of inactivated whole virion vaccine without any adjuvant via an intradermal route on days 0, 7 & 14 days (n+1 doses).

All studies were conducted with an equal number of males and females unless otherwise specified. The control group was injected with saline. Animals were bled from the retro-orbital plexus, 2hours before each immunization on0, 7, 14 & 21 days, and serum was separated and stored at −20°C until further use.

Pooled and individual sera from vaccinated mice and rabbits were used to test the antigen-specific antibody binding titer and antibody isotyping profile by Enzyme-Linked Immunosorbent Assay (ELISA). Pooled or Individual sera from all vaccinated species (mice, rabbits & rats) were used to test neutralization antibody titer by Plaque Reduction Neutralization Test (PRNT_90_) or Micro Neutralization Test (MNT_50_).

### 9. Enzyme-linked immunosorbent assay (ELISA)

ELISA tests were performed as per standard protocols specifically for this project. Microtiter plates were coated with SARS-CoV-2 specific antigens (whole inactivated antigen or spike, S1 /Receptor Binding Domain (RBD)/ nucleocapsid (N) at a concentration of 1μg/ml, 100μl/well in PBS pH 7.4). After incubation, wells were added with Goat Anti-mouse IgG HRP(Santa Cruz Biotechnology, USA) conjugated antibody for mouse sera samples, and Goat anti-rabbit IgG HRP conjugate antibody(Santa Cruz Biotechnology, USA) (dilution 1:2500) for rabbit sera samples and incubated for 1hr at RT. Threshold (Mean + 3SD) was established by taking the absorbance of negative control (PBS) group, or pre-immune sera and antigen-specific endpoint titers were determined. The antibody dilution, at which absorbance is above the threshold, was taken as antigen-specific antibody endpoint titers.

### 10. Immunoglobulin (IgG) Subclass

Th1-dependent IgG2a vs. Th2 -dependent IgG1 antibody subclasses were determined from mice vaccinated sera as previously described ^37^. Briefly, 96 well microtiter plates were coated with various SARS-CoV-2 specific antigens (whole Inactivated antigen, S1, Receptor Binding Domain (RBD), nucleocapsid (N) at a concentration of 1μg/ml, 100μl/well in PBS pH 7.4) and kept at 2-8°C for overnight. The next day, plates were washed with washing buffer (PBST) and blocked with a blocking buffer (PBS with 2% BSA) at RT for one hour. serially diluted (dilutions from 1:50 to 819200 in PBS, 0.1% BSA, 0.05% Tween™20, 0.02% sodium azide) pooled or individual sera from hyperimmunized animals (mice/rabbits) and incubated at 37°C for 2hrs. After incubation, wells were washed and added with anti-mouse IgG1 or IgG2a HRP conjugate antibodies at a dilution 1:2500. After incubation of the plate for 1hr at RT, wells were washed, and 3,3⍰,5,5⍰-tetramethylbenzidine (TMB) was added as a substrate to develop color. Absorbance was read at 450 nm. Threshold (Mean + 3SD) was established by taking the absorbance of negative control (PBS) group, or pre-immune sera and antigen-specific endpoint titers were determined. The antibody dilution, at which absorbance is above the threshold, was taken as antigen-specific antibody endpoint titers.

### 11. Cytokine (IFNγ & IFNα) Estimation by ELISA

To determine IFNγ, Enzyme-Linked Immunosorbent Assay (ELISA) was performed according to the instruction manual. Briefly, the capture antibody was first diluted in coating buffer and added 100 μL to each well in 96-well microplate. Plates were incubated overnight at 2-8 °C. Coated plates were then washed with wash buffer (PBST). After washing, these plates were blocked using 1x assay diluent for 1hr at room temperature followed by washing with PBST. Serial dilutions of Top Standard were prepared to make the standard curve. Similarly, 4-fold dilutions (1:4, 1:16 & 1:64) of serum samples were e prepared and added to wells in triplicates, and the plate was incubated at room temperature for 2hrs. After washing the plate, 100 μL/well of detection antibody diluted in 1X Assay diluent was added and incubated at room temperature for 1hr. Later, 100 μL/well of Avidin-HRP* diluted in 1X Assay diluent was added and incubated at room temperature for 30 minutes. Finally, after washes, 100 μL of substrate solution was added to each well and incubated at RT for 15 minutes. The reaction was stopped by the addition of 50 μL of 2N H_2_SO_4_ to each well, and the plate was read at 450 nm.

PBMCs cell culture supernatant was used to estimate IFNα using The VeriKine Human Interferon Alpha ELISA Kit (PBL Assay Science, USA, Cat log# 41100). The assay was performed as per the manufacturer’s instructions. Briefly, Pre-coated plates were incubated with diluted standard (range 500-12.5 pg/ml) or culture supernatant, for 1hr at room temperature. Later, the diluted antibody and HRP solution were added sequentially. TMB was used as a substrate, followed by the addition of stop solution. The plate was read at 450nm.

### 12. Intracellular Staining

Vaccinated splenocytes (2×10^6^/ml) were cultured in 24 well plates and stimulated with inactivated SARS-COV-2 antigen (1.2 μg/ml) or PMA (25 ng/ml, cat # P8139; Sigma) and Ionomycin (1 μg/ml, cat # I0634, Sigma) along with Protein transport inhibitor (Monensin, 1.3μl/ml cat # 554724, BD biosciences). Cells were washed and centrifuged at 1000rpm for 5-10min and stained with APC-Cy™7 Rat Anti-Mouse CD3 (clone: 17A2, Cat # 560590, BD Biosciences), FITC Rat Anti-Mouse CD4 (Clone: H129.19, Cat # 553650, BD Biosciences), and PE-Cy™7 Rat Anti-Mouse CD8a (Clone: 53-6.7, Cat # 552877, BD Biosciences) for 30 minutes at 4°C. Cells were again washed twice with PBS and fixed using fixation/Permeabilize solution (Cat # 554722, BD Biosciences) for 20 mins at 4°C. Following fixation/permeabilization, cells were washed with 1x permeabilization buffer and stained with intracellular cytokines (IFN-L (BV421 Rat Anti-Mouse IFN-γ, Clone: XMG1.2, cat # 560660, BD Biosciences) for 30 mins at 4°C. Cells were washed and resuspended in 500μl FACS buffer (Cat # 554657, BD Biosciences). All samples were acquired using BD FACSVerse (BD Biosciences).

### 13. Cytokine Estimation

To assess the secretion of Th1 or Th2 mediated cytokines, if any, and to differentiate between Algel1 and Algel2, we used vaccinated mice sera samples collected at various time points (Day 0, 7, 14, 21 & 28, 7 days post-vaccination) and measured Cytokines using the BD CBA Mouse Th1/Th2/Th17 Cytokine Kit (BD Bioscience, San Jose, CA, USA). Sera samples were processed as per the manufacturer’s instructions. Briefly, the kit was used for the simultaneous detection of mouse IL-2, IL-4, IL-6, IFN-γ, TNF, IL-17A, and IL-10 in a single sample. For each sample, 50 μL of the mixed captured beads, 50 μL of the unknown serum sample or standard dilutions, and 50 μL of phycoerythrin (PE) detection reagent were added consecutively to each assay tube and incubated for 2 h at room temperature in the dark. The samples were washed with 1 mL of wash buffer for 5 min and centrifuged. The bead pellet was resuspended in 300 μL buffer after discarding the supernatant. Samples were measured on the BD FACS Verso and analyzed by FCAP Array Software (BD Bioscience).

### 14. Plaque Reduction Neutralization Test (PRNT_90_)

The Plaque reduction neutralization test was performed in a biosafety level 3 facility. To perform PRNT_90_, Vero CCL-81 cell suspension (1.0 × 10^5^ /mL/well) was added in duplicates in 24-well tissue culture plates and cultured in a CO_2_ incubator at 37°C for 16-24 hrs. Vaccinated serum samples were inactivated by keeping in a 56°C-water bath for 30 min. Serial dilutions (4 fold) of vaccinated serum samples were mixed with the virus, which can form 50 plaque-forming units and then incubated for 1 h at 37°C. The virus–serum mixtures were added onto the preformed Vero CCL-81 cell monolayers and incubated 1 h at 37°C in a 5% CO_2_ incubator. The number of plaques was counted, and the Neutralizing antibody titer was determined based on the 90% reduction in the number of plaque count, which was further analyzed using 50% ProbitAnalysis^38^. A neutralization antibody titer < 1:20 considered negative, while that of > 1:20 considered as positive.

### 15. Micro Neutralization assay (MNT)

The serum of the animal to be tested was inactivated in a 56°C -water bath for 30 min. Serum was successively diluted 1:8 to the required concentration by a 2-fold series, and an equal volume of challenge virus solution containing 100 CCID_50_viruses was added. After neutralization in a 37°C incubator for two hours, a 1.0 x 10^5^ /mL cell suspension was added to the wells (0.1 mL/well) and cultured in a CO_2_ incubator at 37⍰ for 3-5 days. The Karber method ^35^by observing the CPE was used to calculate the neutralization endpoint (convert the serum dilution to logarithm), which means that the highest dilution of serum that can protect 50% of cells from infection by challenge with 100 CCID50 virus is the antibody potency of the serum. A neutralization antibody potency < 1:20 is negative, while that R 1:20 is positive.

### 16. Mutagenicity Assay (Bacterial Reverse Mutation)

The mutagenic potential of the Adjuvant, Algel-IMDG, was evaluated by Bacterial Reverse Mutation assay through plate incorporation and pre-incubation methods using *Salmonella typhimurium* strains TA 1535, TA 1537, TA 98, TA 100, and TA 102 following OECD Guidelines for Testing of Chemical^14^, with and without S9. Toxicity was apparent either as a reduction in the number of His+ revertants or as an alteration in the auxotrophic background (*i.e*., background lawn).

### 17. Maximum Tolerated Dose Test or Single Dose Toxicity Study

Two animals (Swiss Albino mice and Wistar Rats) species were tested with Algel-IMDG with a single maximum dose (containing 200μg Algel and 20μg TLR7/8 agonist molecule). Animals (Swiss Albino mice and Wistar Rats) were administered via an intramuscular route with Algel-IMDG on day 0 and observed for clinical signs, mortality, and changes in body weight if any up to 14 days. The site of injection was also observed for erythema and edema at 24, 48, and 72 hours after dosing to detect the local tolerance (local reactogenicity) of Algel-IMDG. All animals were necropsied and examined macroscopically. Histopathology was performed for the site of injection.

### 18. Repeated dose toxicity

Studies were performed following both national and international guidelines in compliance with OECD principles of GLP^14, 15, 37-39^. Three animal models (Mice, Rats & Rabbits) were administered via an intramuscular or intraperitoneal with three doses (N+1) of antigen or adjuvanted vaccine at different concentrations. All animals were observed for mortality during the experimental period. Blood collected on day 2 and 21 from the main groups and day 28 from the recovery group were analyzed for detailed clinical pathology investigations. Animals were euthanized either on day 21 (main groups) or on day 28 (recovery groups) and necropsied, and organs were evaluated for macroscopic and microscopic findings.

#### Test system

The test system *viz*., Swiss albino mice (SA), BALB/c mice, Wistar rats, and New Zealand White (NZW) rabbits (*in vivo* models) were sourced from CPCSEA approved vendor and strains of *Salmonella typhimurium* (Moltox, Switzerland) for *in vitro* assay, and these test systems were selected as per the recommendations of WHO guidelines ^16, 39^ and Schedule Y (2019)^15^. The studies were conducted in an equal number of adult males and females except in the BALB/c mice study, where only females were used. The control group was administered with PBS.

#### Treatment regimen

The adjuvanted vaccines or adjuvants alone were administered intramuscularly (IM) in quadriceps muscles of the hindlimb on days 0, 7, and 14 (n+1) with full Human single dose (HSD) to NZW rabbits and SA mice and higher dose than HSD to Wistar rats and full HSD to. In BALB/c Mice, 1/20^th^ HSD was administered intraperitoneally. The animals were observed up to 14 days, post last dose.

#### Experimental Design - Adjuvant alone

Maximum Tolerated Dose (MTD) studies were conducted using Wistar rats and Swiss Albino mice with ten animals in each study. The animals were treated with a single dose of Algel-IMDG at the dose of 200 μg /animal and observed for 14 days. Two repeated dose toxicity studies with Algel and Algel-IMDG in Wistar rats and Swiss Albino mice were performed. Control and reversal groups were maintained. The site of injection was observed for erythema and edema at 24, 48, and 72 hours after dosing to detect the local tolerance (local reactogenicity) of Algel-IMDG. All animals were necropsied and examined macroscopically. Histopathology was performed for the site of injection.

#### Experimental Design - Adjuvanted Vaccines

Four repeated dose toxicity studies were performed with Adjuvanted vaccines in Wistar Rats, New Zealand White Rabbits, BALB/c Mice, and Swiss Albino Mice.

Algel alone, Antigen alone, Adjuvanted Vaccine with Algel, and adjuvanted vaccine with Algel-IMDG along with control and recovery groups were assigned. We have tested adjuvants in the highest concentration of 300ug and antigen at the concentration of 9ug, to evaluate safety.

#### In-life Observations

All animals were observed twice daily for mortality. Clinical signs were recorded twice a day from day 0 to 2 and once daily thereafter. The cage side observations included changes in the skin, fur, eyes, and mucous membranes and clinical signs observed for edema, erythema, alopecia, irritation, necrosis, locomotor activity, lacrimation, hyperthermia, and hypothermia, etc. The body weight of each animal was recorded once daily after the first dose for a week and weekly once thereafter. Mean body weights and mean body weight gain was calculated for the corresponding intervals. The amount of feed consumed by each cage of animals was recorded once daily after the first dose for a week and weekly once thereafter. Body temperature was recorded for rats and rabbits on day 0, 3 hours, and 24 hours after each dose, and on the day of sacrifice

#### Clinical Pathology Investigations

Detailed clinical pathology was performed using automated equipment as per referred guidelines following validated procedures^145, 37-39^. Blood and urine samples were collected for clinical evaluations (hematology, coagulation parameters, acute phase proteins, serum chemistry, and urinalysis) from all the groups.

#### Necropsy, Organ Weight and Histopathology

Animals were euthanized by carbon dioxide asphyxiation and necropsied. Organs, as per WHO guidelines, which included spleen, thymus, and draining lymph nodes (inguinal for IM), were collected from all terminally sacrificed animals, and macroscopic abnormalities were recorded. Wet weights for organs such as brain, thymus, spleen, ovaries, uterus, heart, kidneys, testes, liver, adrenals, lungs, epididymides, and prostate with seminal vesicles and coagulating glands were recorded.

### 18. Statistical Methods

Statistical Analysis was performed in R 4.0.1. We used two-sided one sample t-test with 5% level of significance for continuous variables which followed a normal distribution. To test the significance of the sample, mean and for the variables that do not satisfy the normality assumption, we used the Mann-Whitney test with 5% level of significance to test the significance of median.

## Acknowledgments

We express our sincere gratitude to Dr. Rudragouda Channappanavar, University of Tennessee) for the scientific review of this paper. Dr. Zaiham Rizvi from the Translational Health Sciences and Technology Institute, Faridabad, India, assisted with cell-mediated response analysis. Our special thanks to Dr. Sunil David (ViroVax LLC) for giving us adjuvant samples for evaluation in the initial development phase of the project. This vaccine candidate could not have been developed without the efforts of Bharat Biotech’s manufacturing and quality control teams. All authors would like to express their gratitude for all the frontline health care workers during this pandemic.

## Author Contributions

All listed authors meet the criteria for authorship set forth by the International Committee for Medical Editors and have no conflicts to disclose. BG., J.H., S.R., J.J., led the immunogenicity and safety preclinical experiments. H.J., V.D., N.M., V.K.S., S.P., K.M.V the manufacturing and quality control efforts. KMV, P.S., and E.R. provided technical assistance with design, analysis, and manuscript preparation. Y.P., S.G., S.A., M.S., A.B., A.P., B.B., N.G of ICMR-NIV, Pune conducted electron microscopy and neutralizing antibody assays. A.A conducted cell-mediated response related assay activities at THSTI. J.J., R.R., led the safety assessments in animals.

