## Supplemental Figures for "Evaluation of Safety and Immunogenicity of an Adjuvanted, TH-1 Skewed, Whole Virion InactivatedSARS-CoV-2 Vaccine - BBV152"

**Figure S1: Mutagenicity dose-response curve (with and without metabolic activation)**

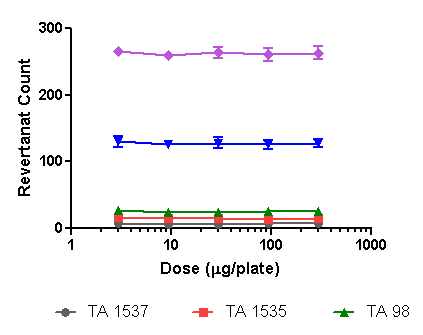

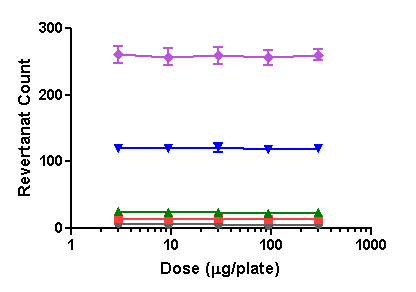

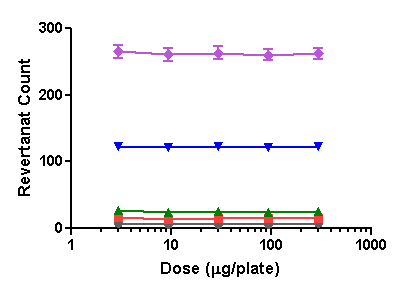

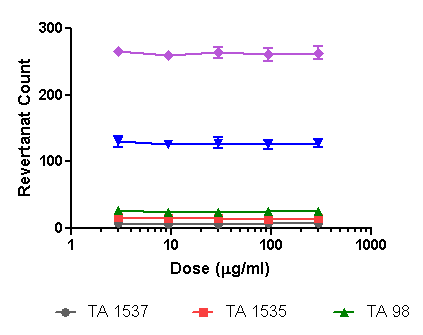

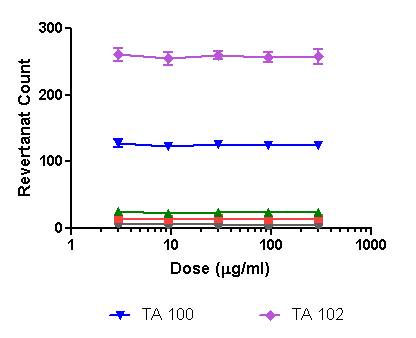

**(+S9)**

**(-S9)**

**A. Plate Incorporation Method**

**B. Pre-Incubation Method**

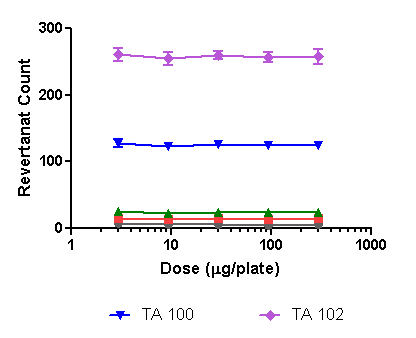

*Revertant colonies observed when five strains (TA 1537, TA 1535, TA 98, TA 100, TA 102) were treated with Algel-IMDG, in the presence and the absence of S9, by two methods; A. Plate Incorporation method; B. Pre-Incubation method.*

**Figure S2: Percent Body Weight Gain in Wistar Rats when administered.**

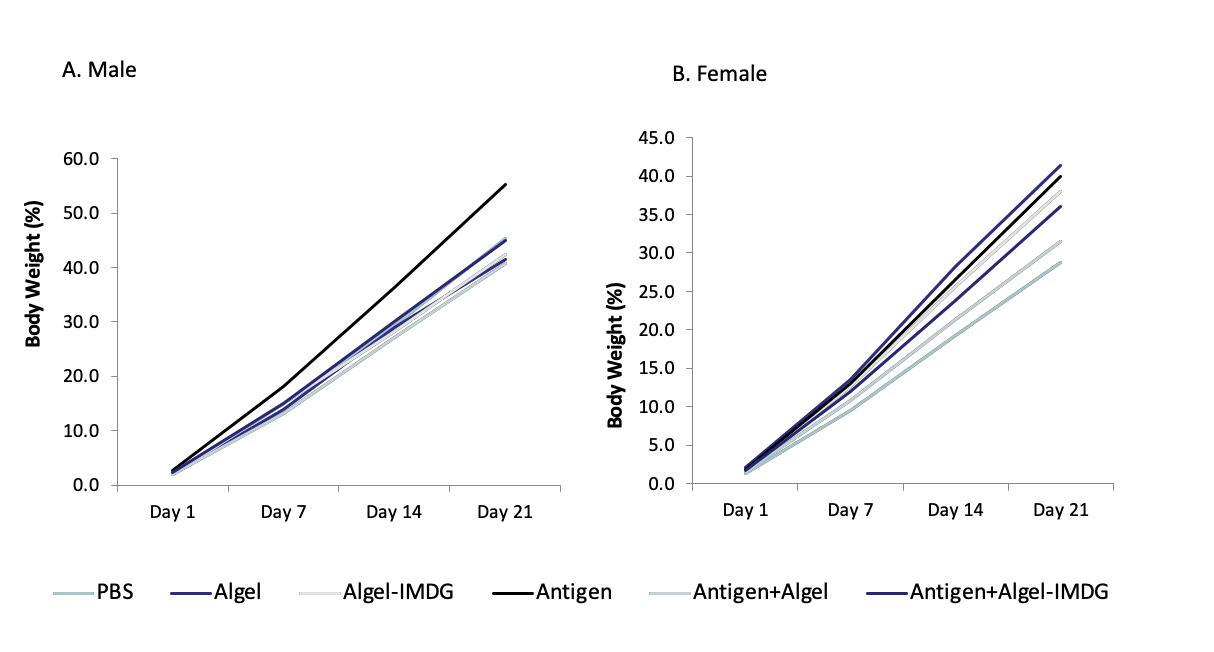

*Percent body weight recorder during the experimental period (Day 1, 7, 14, and 21). Groups received any of the following: Phosphate buffer saline (PBS), Algel, Algel-IMDG, Antigen, Antigen+Algel, Antigen+Algel-IMDG. A. Male; B. Female Percent Body Weight Gain in Males & Females Monitored to Evaluate the Safety in Wistar Rats.*

**Figure S3: Safety evaluation of Haematology parameters in vaccinated Wistar Rats.**

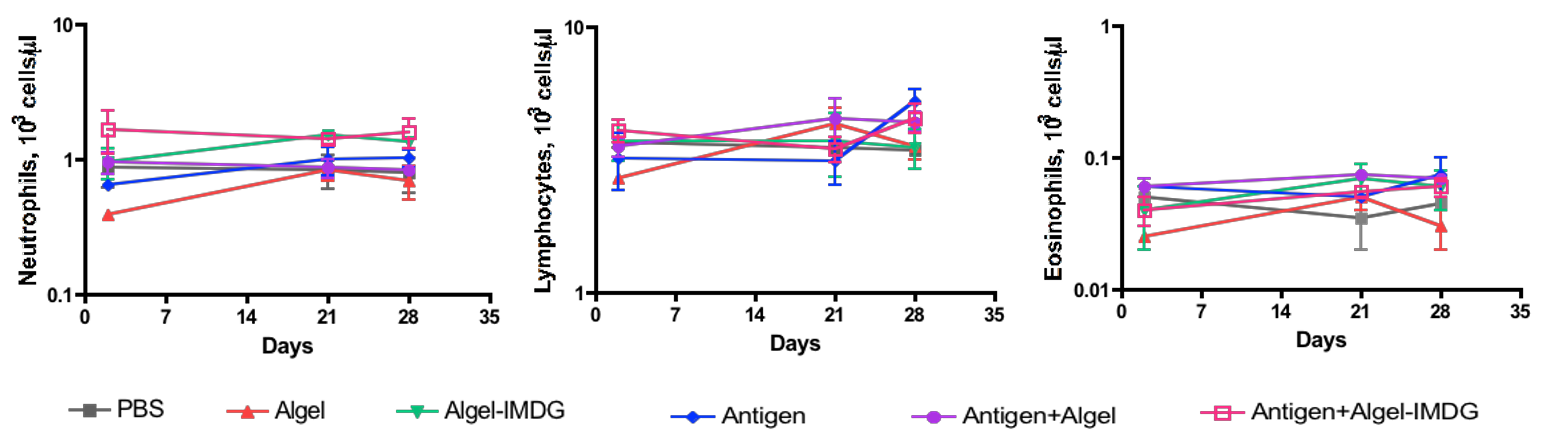

*Representative haemotology parameters such as neutrophils, lymphocytes, eosinophils were evaluated on day 2, 21, and 28 of phosphate buffer saline (PBS) or antigen+ Algel-IMDG. Error bars signify means ± S.D.*

**Figure S4: Algel-IMDG induces more Macrophage infiltration at the site of injection (quadriceps muscles).**

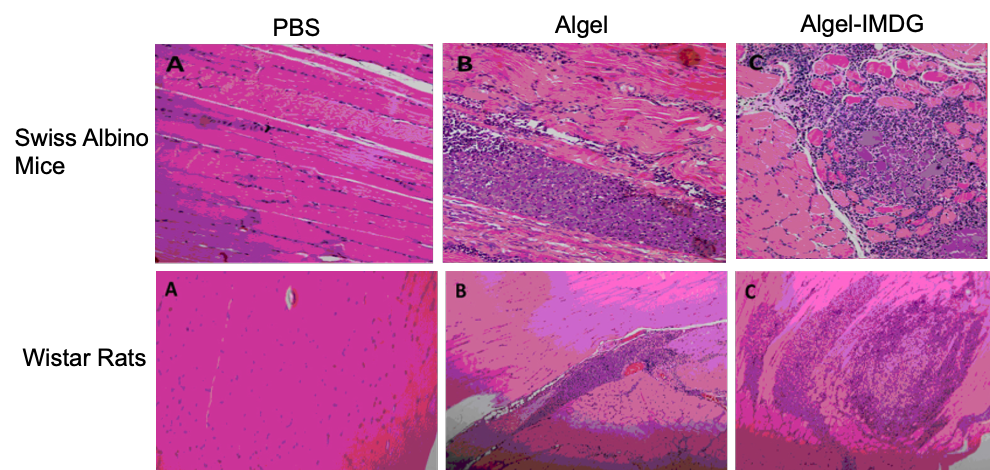

*Representative photomicrographs of the site of injection (quadriceps muscles) from Swiss Albino mice (top) and Wistar rats (bottom) stained with hemotoxylin and eosin, original magnification was 40X.* ***A****. Phosphate Buffer Saline;* ***B****. Algel (300 µg) showing infiltration of macrophages containing bluish stained material;* ***C****. Algel-IMDG (300 µg) showing infiltration of macrophages containing bluish stained material.*

**Figure S5: Adjuvanted Vaccines Induces Macrophage Infiltration *at The Site of Injection***

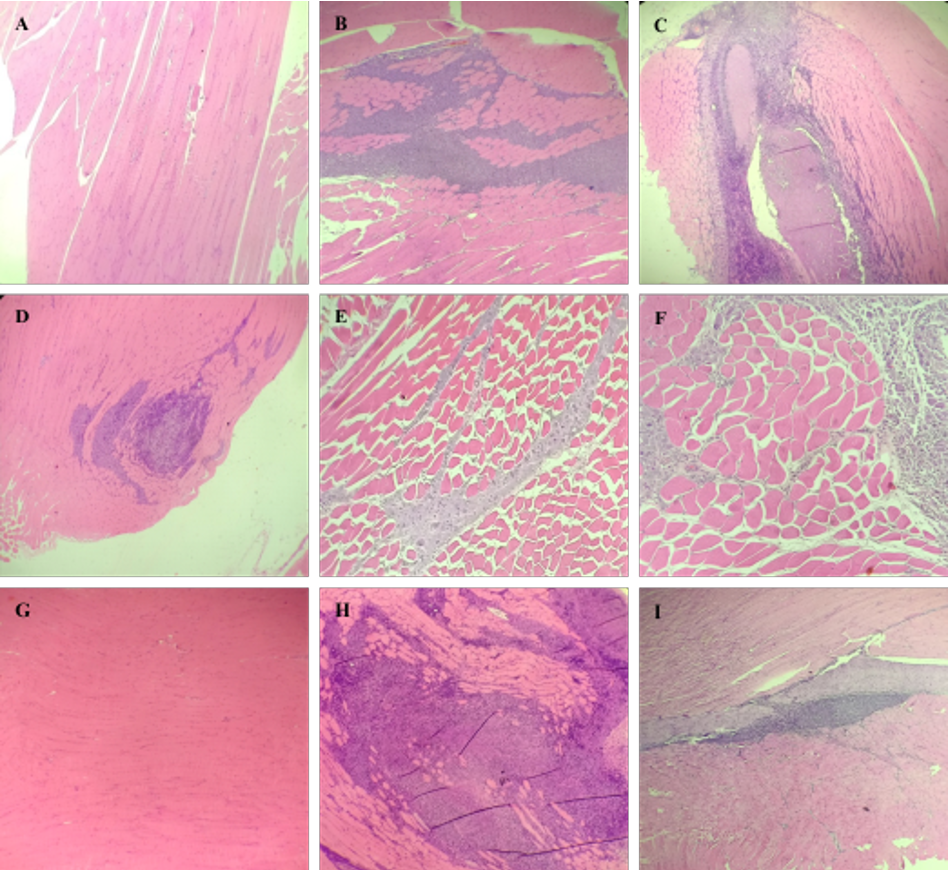

***Top Row: Representative photomicrographs of the site of injection (quadriceps muscles) from Wistar rats.*** *Animals vaccinated with (A) Phosphate buffer Saline, (B) Antigen (9µg) + Algel (300µg) group showed macrophage infiltration of macrophages with a bluish stained material, (C) Antigen (9µg) + Algel-IMDG (300µg) group with chronic inflammation around the site of injection, (D) Recovery group Antigen (9µg) + Algel-IMDG (300µg) showing infiltration of macrophages. (hematoxylin and eosin, original magnification was 40X).*

***Middle Row: Representative photomicrographs of the site of injection (quadriceps muscles) from New Zealand White Rabbits.*** *Animals treated with (E) Antigen (6 µg) + Algel (250 µg) (F) Antigen (6 µg) + Algel-IMDG (250 µg) showed macrophage infiltration with a bluish stained material (hematoxylin and eosin, original magnification was 40X).*

***Bottom Row: Representative photomicrographs of the site of injection (quadriceps muscles) from Swiss Albino mice.*** *Animals vaccinated with (G) Phosphate buffer Saline, (H) Antigen (6 µg) + Algel-IMDG (250 µg) group showing chronic inflammation around test item deposits, (I) Recovery group Antigen (6 µg) + Algel-IMDG (250 µg) showing infiltration of macrophages. (hematoxylin and eosin, original magnification was 40X).*

**Figure S6: Photomicrographs of tissues of Wistar Rats**

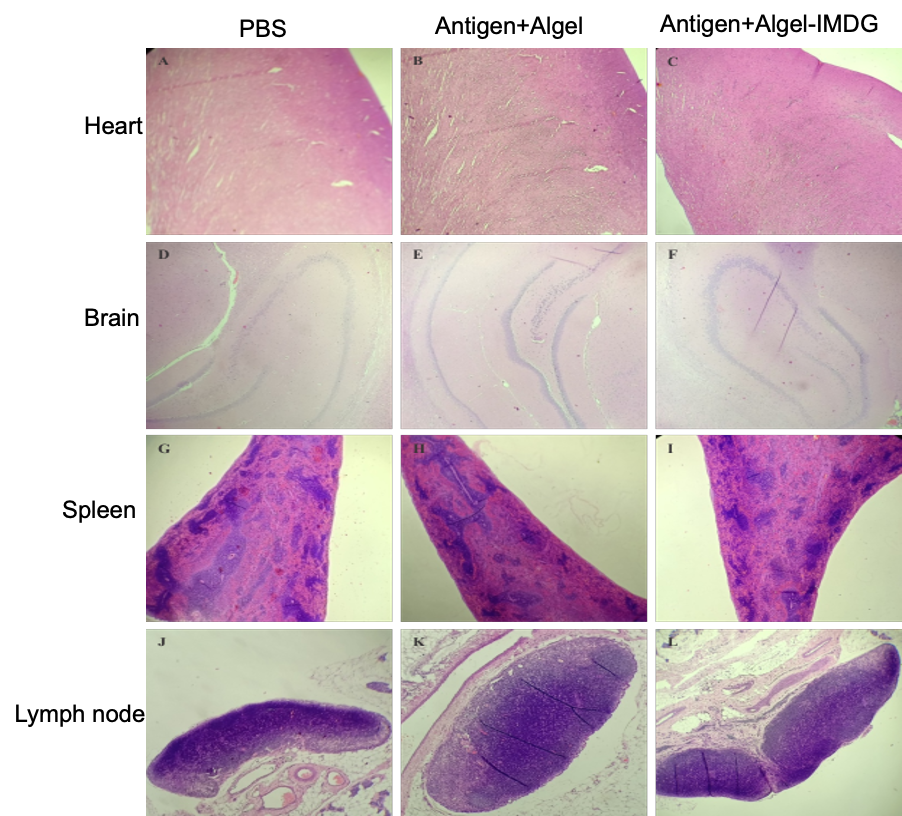

*Representative photomicrographs of Heart, Brain, Spleen and Inguinal lymph nodes from Wistar Rats. These organs were within normal histological limits. (A) Heart - Control, (B) Heart - Antigen 9µg + Algel-1 300µg, (C) Heart - Antigen 9µg + Algel-2 300µg, (D) Brain - Control, (E) Brain - Antigen 9µg + Algel-1 300µg, (F) Brain - Antigen 9µg + Algel-2 300µg, (G) Spleen - Control, (H) Spleen - Antigen 9µg + Algel-1 300µg, (I) Spleen - Antigen 9µg + Algel-2 300µg, (J) Inguinal lymph nodes - Control, (K) Inguinal lymph nodes - Antigen 9µg + Algel-1 300µg, (L) Inguinal lymph nodes - Antigen 9µg + Algel-2 300µg (hematoxylin and eosin, original magnification was 40X).*

**C**

**Figure S7: Representative photomicrographs of Liver, Kidney, and Lungs from New Zealand White Rabbits.**

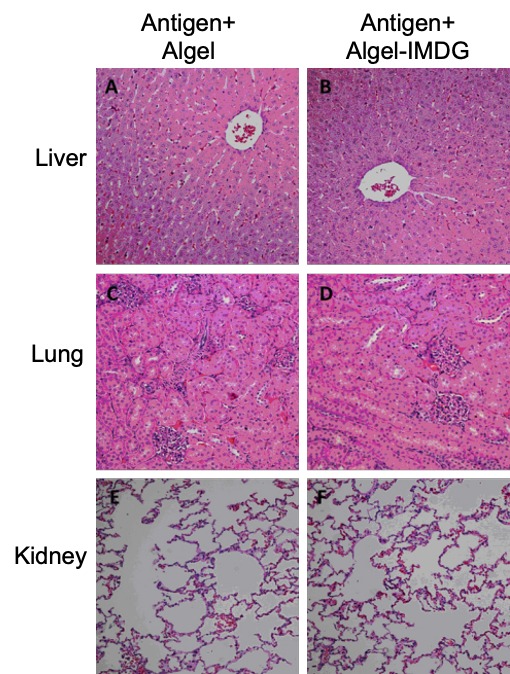

*Representative photomicrographs of Liver, Kidneys & Lungs from rabbits, when administered with adjuvanted vaccines (BBV152B&C). Histopathological sections of all Organs showed within normal limits: Concentration of antigen used in both adjuvanted formulation was 6μg. Concentrations of adjuvant were: Algel (300μg) and Algel-IMDG (300μg).*

**Figure S8: Comparison of the immune response of the adjuvanted vaccine in BALB/c mice at 7-day and 14-day dose intervals.**

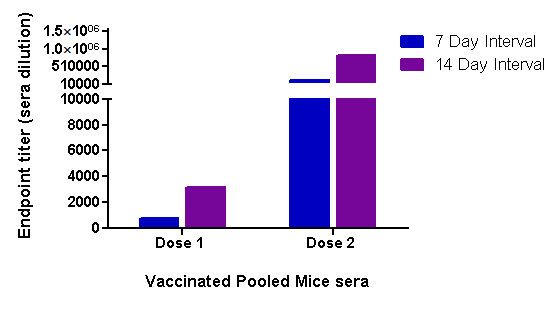

*Balb/C mice (n=10) were administered with adjuvanted vaccine formulation (BBV152B) via IM route with Full Human Single Dose (HSD): S1 specific Total IgG antibody binding titer performed by ELISA post-immunization.*

**Figure S9: Comparison of the immune response of antigen alone with the adjuvanted vaccine in BALB/c mice.**

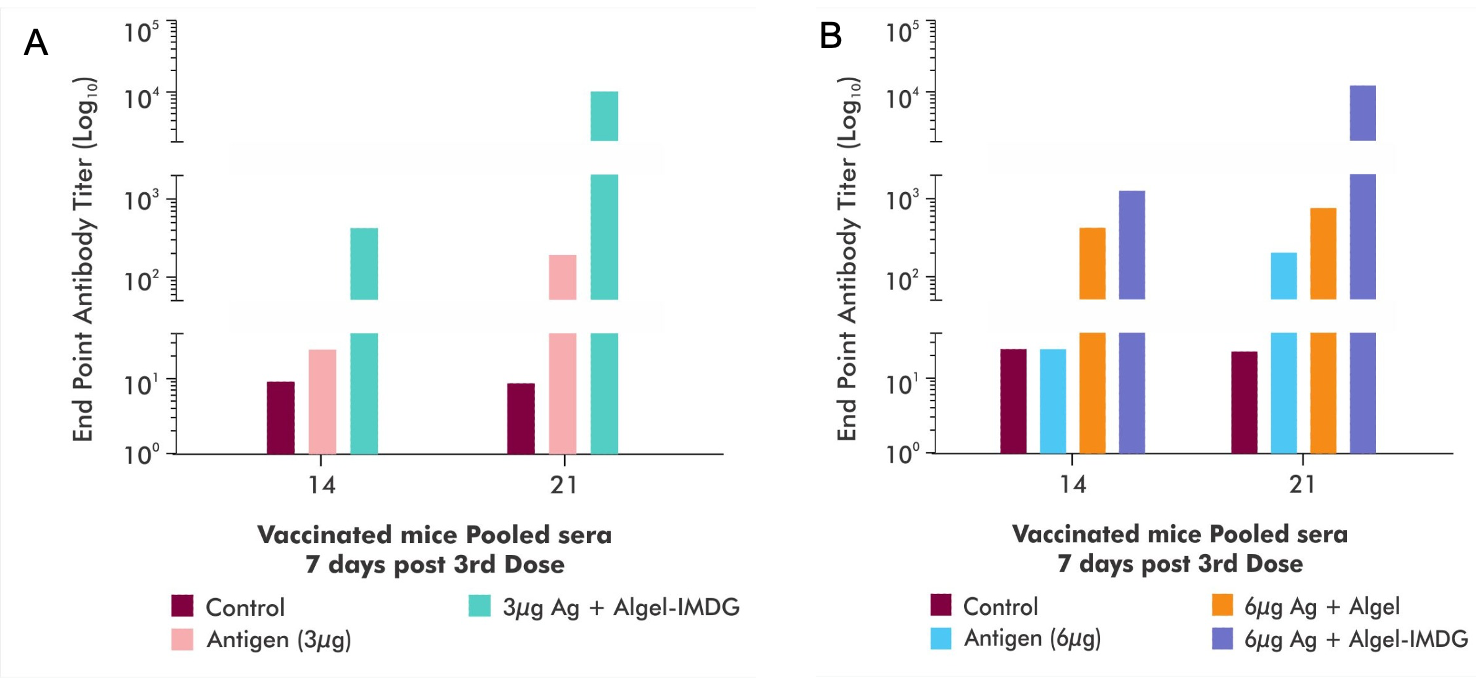

*Balb/C mice (n=10) were administered with antigen alone and adjuvanted vaccine formulations via IP route, either with 1/20^th^ Human Single Dose (HSD): S1 specific Total IgG antibody binding titer performed by ELISA, using sera collected at various time points (Day 14 & 21). A. 3μg of antigen alone and adjuvanted vaccine; B. 6 μg of antigen alone and adjuvanted vaccine.*
